# Full activation of an α7-α3β4 nicotinic receptor complex avoids receptor desensitization in human chromaffin cells

**DOI:** 10.1101/2020.11.11.360537

**Authors:** Amanda Jiménez-Pompa, Sara Sanz-Lázaro, José Medina-Polo, Carmen González-Enguita, Jesús Blázquez, J. Michael McIntosh, Almudena Albillos

## Abstract

α7 nicotinic receptors have been involved in numerous pathologies. A hallmark of these receptors is their extremely fast desensitization, a process not fully understood yet. Here we show that human native α7 and α3β4 nicotinic receptors physically interact in human chromaffin cells of adrenal glands. The full activation of this α7-α3β4 receptor complex avoids subtypes receptor desensitization, leading to gradual increase of currents with successive acetylcholine pulses. Instead, full and partial activation with choline of α7 and α3β4 subtypes, respectively, of this linked receptor leads to α7 receptor desensitization. Therefore choline, a product of the acetylcholine hydrolysis, acts as a brake by limiting the increase of currents by acetylcholine. Very importantly, the efficiency of the α7-α3β4 interaction diminishes in subjets older than 50 years, accordingly increasing receptor desensitization and decreasing nicotinic currents. These results open a new line of research to achieve improved therapeutic treatments for nicotinic receptors related diseases.

## Introduction

Nicotinic acetylcholine receptors are ligand-gated cationic channels that mediate fast synaptic transmission. There are different nicotinic receptor subunits, α1–7, α9, α10, β1-β4, γ, δ, and ε in mammals, which assemble into pentamers forming different heteromeric and homomeric subtypes. In particular, the α7 subtype forms homomeric receptors that exhibit very fast desensitization with repetitive pulses. However, heteromeric α7β2 receptors have also been reported, and show similar pharmacological sensitivities compared to the homomeric receptor. α7 receptors are expressed together with other receptor subtypes in neuronal and neuroendocrine cells. Whether these receptor subtypes interact with each other is unknown. A previous report of Criado and colleagues (Criado *et al.*, 2012) showed that α7 subunits possess the capacity to stably associate with β4 or α3 subunits expressed in *Xenopus oocytes*. Here we pose the hypothesis that a physical interaction, and a mutual regulation of expression and functional activity exists between α7 and non-α7 receptor subtypes.

We evaluated that hypothesis in a human native system, the chromaffin cells of human adrenal glands obtained from organ donors. Human chromaffin cells of the adrenal gland are modified postganglionic sympathetic neurons. These cells are the major adrenaline-secreting cells in our organism. They also release a cocktail of compounds such as opioids, ATP or calcium to the bloodstream by exocytosis under stressful situations such as the flight or fight response. Chromaffin cells allow the study of these receptor subtypes and their involvement in neurotransmitter secretion process in an incomparable manner (Albillos *et al.*, 1997; Perez-Alvarez & Albillos 2007). In humans, these cells mainly express α7 and α3β4 receptor subtypes (Hone *et al.*, 2015; Hone *et al.*, 2017; Perez-Alvarez *et al.*, 2012a; Perez-Alvarez *et al.*, 2012b). In this study we have discovered key aspects for the understanding the functioning of nicotinic receptors. These findings include the physical interaction between nicotinic receptor subtypes. One of the most relevant consequences of it is that the α7 receptor does not desensitize when the α3β4 subtype is simultaneously and fully activated. The interaction between receptor subtypes decreases with age, and this causes an increase in the rate of desensitization of α7 nicotinic currents. In addition, the expression of receptor subtypes is larger in men with respect to women. These data open a new line of research in the field of nicotinic receptors.

## Results

### Repetitive pulses of ACh elicit an increase of currents

Repetitive pulses of acetylcholine (ACh) (300 μM, 500 ms, applied every 90 s) elicited currents of increasing peak amplitude and charge (Fig. 1A). Initial peak current and charge values amounted to 813.1±103.4 pA and 892.2±176 pC (n=29), respectively. After the application of ACh pulses, the maximal ACh peak current and charge increased to 1638±167.8 pA (****p<0.0001) and 1976±318.7 pC (****p<0.0001) (n=29), respectively, which represents 2-fold and 2.2-fold of increment, respectively (Fig. 1B). The elicited-ACh currents exhibited faster activation kinetics with repetitive pulses (Fig. 1C), decreasing their τ of activation (τ_activation_) from 0.14 ms (n=20) to 0.04 ms (n=11, r Pearson= −0.87, **p<0.01) (Fig. 1D). The number of ACh pulses to reach the maximal increase of current was 7.1±1.1 (n=29). In 1 out 29 cells the increase of current did not occur.

**Figure 1.**
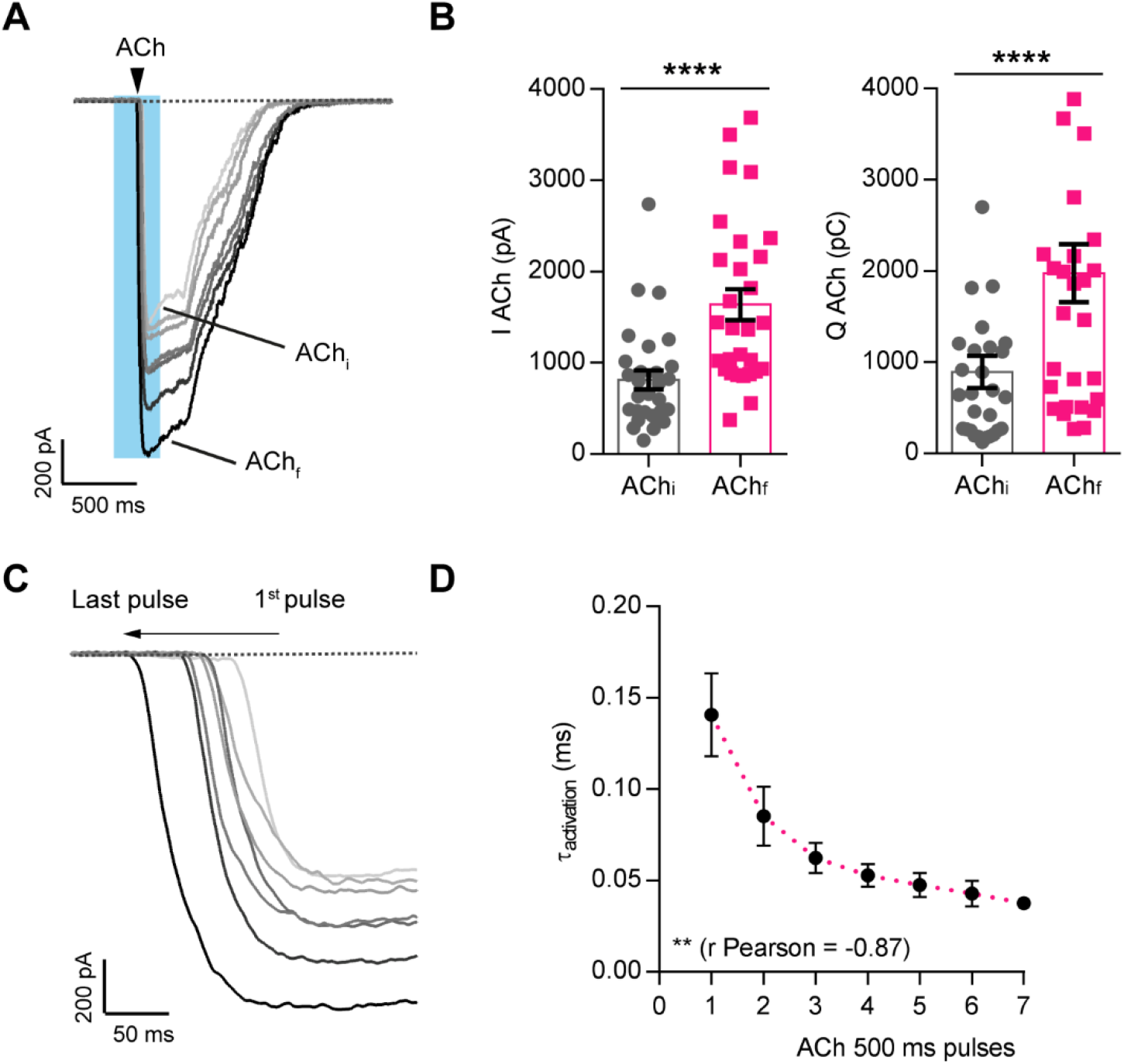
Repetitive pulses of ACh elicit an increase of currents. A) Representative currents elicited by repetitive pulses of ACh (300 μM, 500 ms, applied every 90 s) of increasing peak amplitude and charge. The first and the last current traces elicited by ACh were labeled as AChi and AChf. B) Plotted data of the peak (n=35) and charge (n=29) currents evoked by ACh at the first pulse and at the last pulse when currents reach the maximal value (****p<0.0001). Wilcoxon’s test was performed. C) The shaded part of panel A was enlarged to see in detail how currents are getting faster with repetitive pulses. D) Time constant of activation (τ_activation_) average of the nicotinic currents recorded after application of 7 pulses (number of pulses to reach the maximal ACh peak current) of ACh (n=4-20).

### α3β4 receptors require α7 receptor activation and calcium to increase their activity

In an additional set of experiments we found that the increases of peak current and charge achieved with repetitive ACh pulses were abolished or decreased: i) if the α7 nicotinic current was desensitized previously at the beginning of the experiment by applying three pulses of 3 mM choline, ii) if cells were perfused with the selective α7 receptor blocker ArIB (Whiteaker *et al.*, 2007) at 100 nM during 5 min before the application of ACh pulses and then during the whole experiment, and iii) if ACh was perfused in the absence of calcium. The time courses of the ACh peak current with the corresponding representative recordings under those different conditions are drawn in Fig. 2 (panels B-D). The r Pearson’s correlation coefficients of the nicotinic peak currents achieved were 0.97 (****p<0.0001), −0.98 (****p<0.0001), −0.92 (**p<0.01) and 0.9 (***p<0.001) for control, desensitized α7 receptor, ArIB and free calcium conditions, respectively. Thus, the increases of the current peak and charge achieved with repetitive ACh pulses in control conditions were due to: i) α7 receptor activation, since the observed changes were decreased or abolished in α7 desensitized receptors or ArIB-treated cells; ii) α3β4 receptor activation, since not only the peak current but also the rest of the current were increased with the repetitive stimulation; iii) calcium, since in its absence those variations were diminished. On the other hand, the τ_activation_ of the nicotinic currents elicited with the successive ACh pulses in panels B to D are shown in Fig. 2E-G. The r Pearson correlation coefficients were 0.94 (**p<0.01), 0.45 (no statistical significant differences) and −0.84 (*p<0.05), respectively, for the desensitized α7 receptor, ArIB and free calcium conditions, respectively. Therefore, the decrease of the rate of activation observed in the control situation after repetitive ACh stimulation was mainly due to the recruitment of fast α7 receptors, because the τ_activation_ was increased to a larger extent by previous desensitization of those receptors or by blocking them with ArIB. The removal of calcium slightly increased the τ_activation_, probably because most of the current is carried through the α3β4 receptor subtype in these cells (Hone *et al.*, 2015). In conclusion, α3β4 receptors require α7 receptor activation and calcium to increase their activity.

**Figure 2.**
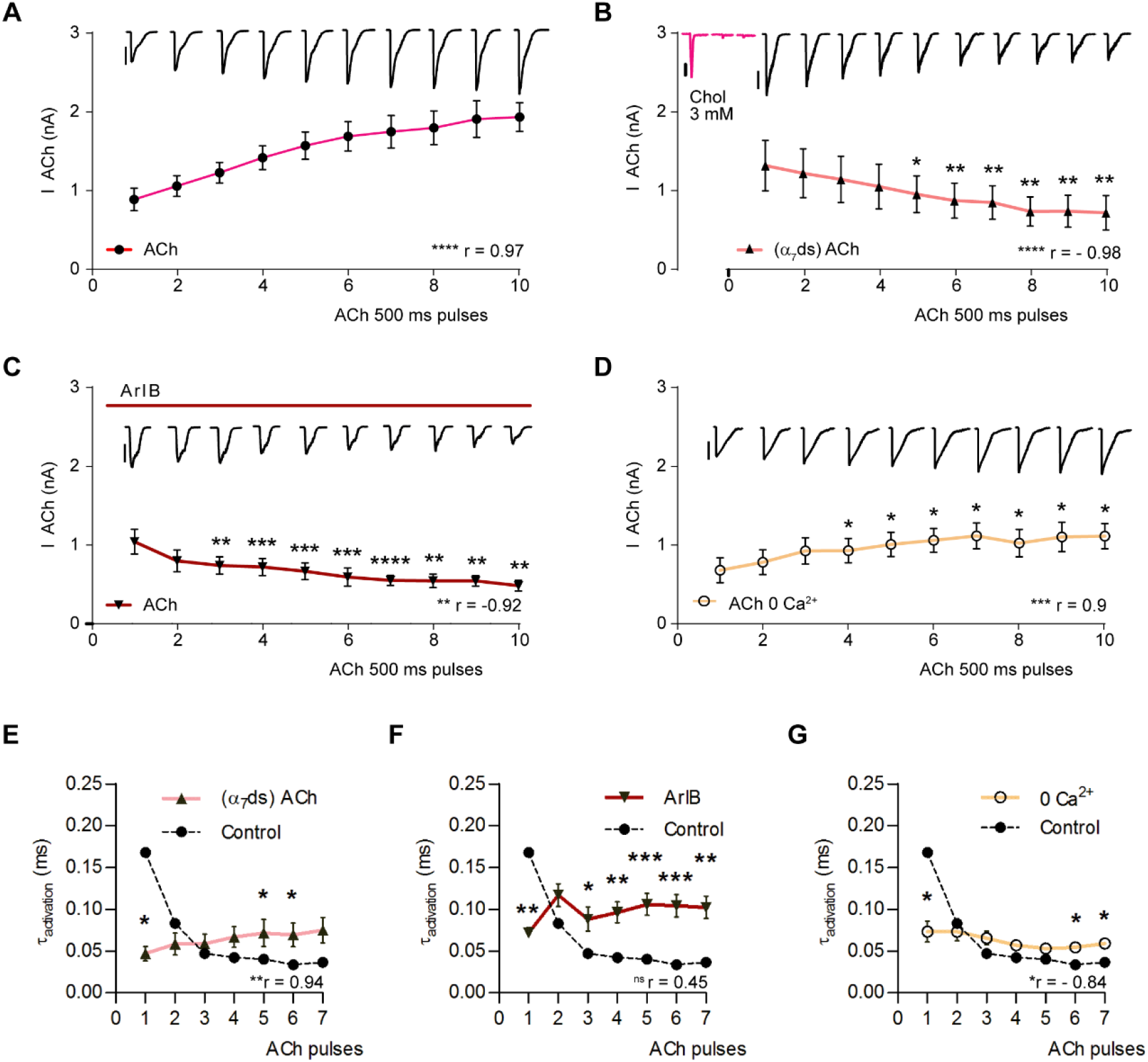
α3β4 receptors require α7 receptor activation and calcium to increase their activity. The ACh peak current is drawn with respect to the number of pulses under different conditions. The corresponding representative recordings for each pulse are shown in the upper part of the panel. A) Control conditions (n=8-12). B) Desensitized α7 receptor: this receptor subtype was desensitized previously to the start of the experiment by applying three pulses of 3 mM choline every 90 s (n=7-10). C) Perfusion with 100 nM ArIB during 5 min before the application of ACh pulses and then during the whole experiment (n=4-12). D) Absence of calcium (n=5-10). Unpaired t-test between each time point and its equivalent point in control conditions. ACh scale: 500 pA. Choline scale: 100 pA. E-G) Time constants of activation of the nicotinic currents from cells of panels A-D under the desensitized α7 receptor (E), ArIB (F) and calcium free (G) conditions, in the 7 successive ACh pulses.

### α7 receptors require α3β4 receptor full activation to avoid their desensitization and to increase their activity

The α7 receptor has been well-characterized as a fast desensitizing receptor. However, here we show that repetitive stimulation with ACh does not desensitize α7 receptors if full α3β4 receptors activation simultaneously takes place. When three pulses of 3 mM choline, an α7 receptor agonist that also activates non-α7 receptors in these cells (Perez-Alvarez *et al.*, 2012b), were applied successively every 90 s, a significant decrease of the choline peak current occurred. The first peak of the current triggered by choline is due to α7 receptor stimulation, while the slow current activated after it is due to partial activation of non-α7 receptors (Hone *et al.*, 2015; Perez-Alvarez *et al.*, 2012a). In these experiments, an initial peak current value of 164.2±38.4 pA was decreased to 60.3±12.6 pA. This corresponds to a decrease of 51.5±13.8% in 11 cells tested (*p<0.05) (Fig. 3A). However, if after the first choline pulse successive pulses of ACh (potentiation protocol) were applied to fully activate α7 and α3β4 receptor subtypes, and then another choline pulse was applied again, the choline elicited peak current this time increased (α7 receptor potentiated currents) (Fig. 3B). On average, peak currents increased from 242.2±78.7 pA to 689.5±119.1 pA after the ACh pulses.

**Figure 3.**
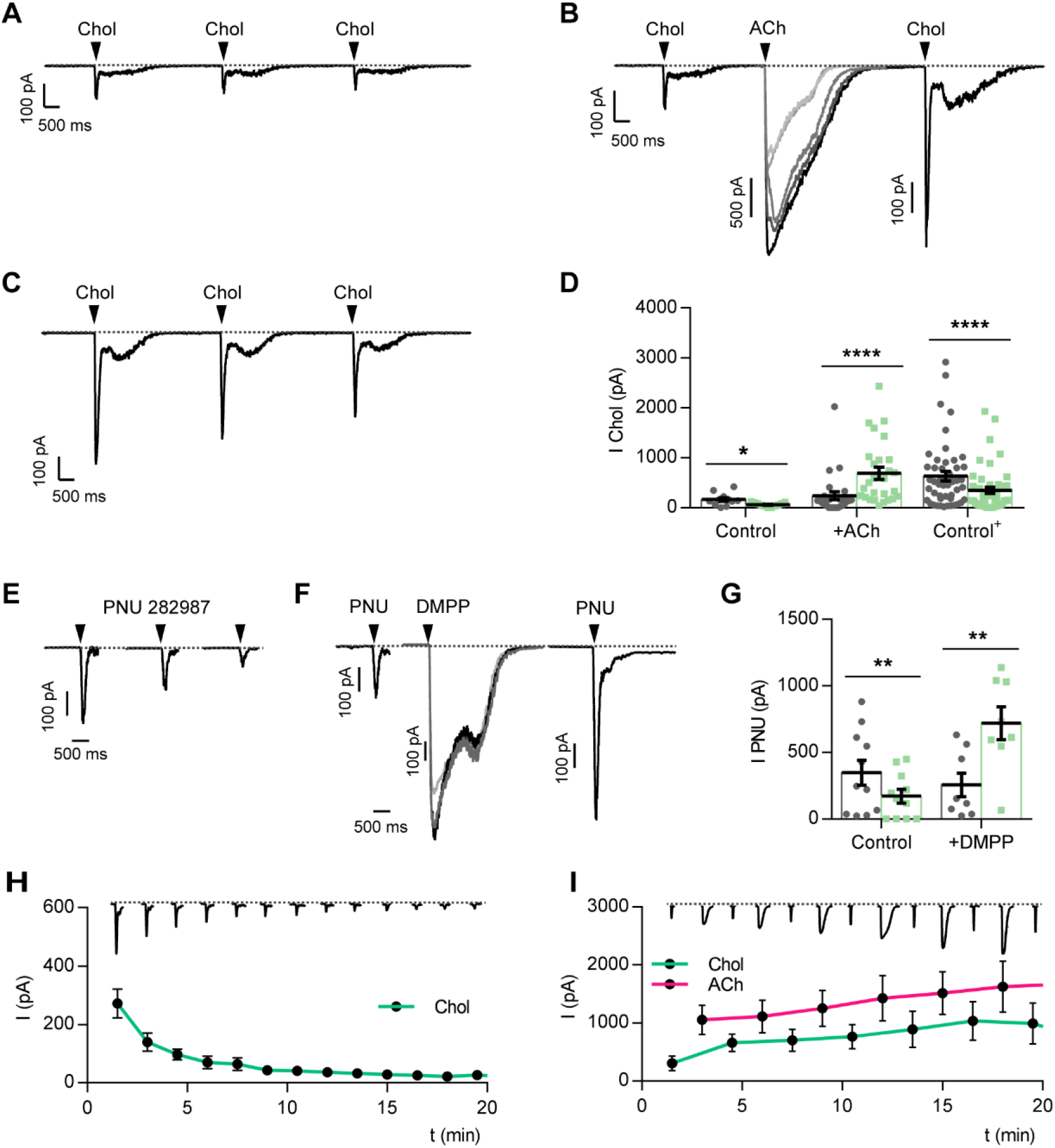
α7 receptors require α3β4 receptor activation to increase their activity. A) Representative recordings of three pulses of 3 mM choline applied every 90 s. B) First a pulse of choline was applied, then pulses of ACh were applied to fully activate α7 and α3β4 receptor subtypes, and later on another choline pulse was applied again, obtaining larger choline-elicited currents (α7 receptor potentiated currents). C) Once the α7 receptor potentiated currents were obtained, another two pulses of choline were applied every 90 s to observe whether the α7 receptor current is desensitized in the third pulse as it happened in panel A. D) A summary of the peak of the choline current elicited in the first (dark grey dots) and the third pulse (green dots) is drawn under control conditions (Control), after the potentiation protocol (+ACh) and after desensitization of the α7 receptor potentiated currents (Control+) (n=11-48). Paired t-test. E-G) the same as in A, B and D), but with PNU282987 instead of choline and DMPP 100 μM instead of ACh (n=8-11). Paired t-test. H and I) Time course of the choline and ACh elicited peak currents obtained with the different pulses. Representative recordings are drawn in the upper part of the figure. The scale of the recordings and the time course is the same (n=6-15).

This meant a 2.8-fold increase in the 27 cells tested (****p<0.0001). The α7 receptor potentiated currents then decreased from 636.3±94.4 pA to 347.3±63.3 pA when three pulses of choline were consecutively applied. Now, the choline elicited currents decreased by 45.3±4.8% (n=48) (****p<0.0001) (Fig. 3C). Data are summarized in the dot plot of Fig. 3D. In order to search whether the same process occurs with a different α7 receptor agonist, 30 μM PNU282987 was perfused instead of choline, and the same protocol was performed. Three sequential PNU282987 pulses caused a decrease of the peak current of 66.6±8.1%, from 348.3±93.3 pA to 172.2±51.4 pA (n=11) (**p<0.01) (Fig. 3E). This decrease of current could be prevented if pulses of the nicotinic agonist DMPP (100 μM) were applied until the maximal current was achieved (5.2±0.8 pulses). PNU282987-elicited peak currents increased from 256.8±88.4 pA to 720.8±123.8 pA. This corresponds to a 2.8-fold increase of the α7 receptor potentiated currents (n=8) (**p<0.01) (Fig. 3F). Data are summarized in the dot plot of Fig. 3G. Thus, the α7 receptor subtype requires full α3β4 receptor activation to avoid its desensitization.

Interestingly, the potentiation of the α7 receptor currents was a very rapid process since a single pulse of ACh was sufficient to avoid the α7 receptor desensitization and to induce the increase of the α7 current. Thirteen choline pulses decreased the current from 280.7±51.6 pA to 27±14 (n=2-15) (90% decrease) (Fig. 3H). The r Spearman correlation was −0.98 (****p<0.0001). However, when 7 pulses of choline were applied alternatively to the ACh pulses, the choline current increased its amplitude by 3-fold from 305.8±127 pA (n=6) to 992.3±354.8 pA (n=3). The ACh elicited current also increased from 1057.6±255.7 pA (n=6) to 1626.6±441.3 pA (1.5-fold increase) (n=3) (Fig. 3I). The r Spearman correlation between ACh and choline currents was 0.69 (****p<0.0001). These results reinforce the idea of the mutual requirement of full activation of α7 and α3β4 receptor subtypes to increase their activity.

### α7 and α3β4 receptors interact physically with each other

The functional interaction above described for α7 and α3β4 receptors might come from a physical interaction. This was investigated in the present study in human chromaffin cells by Förster Resonance Energy Transfer (FRET). Original fluorescence images are shown in Fig. 4A. Cells were treated with the β4* and the α7 receptor markers (asterisk denotes the possible presence of other subunits) BuIA-Cy3 (acceptor) and BgTx-488 (donor), respectively. Under control conditions, FRET efficiency was high, amounting to 22.6±1.5 (n=37) (Fig 4B) which means that, in human chromaffin cells, α7 and α3β4 are physically interacting.

**Figure 4.**
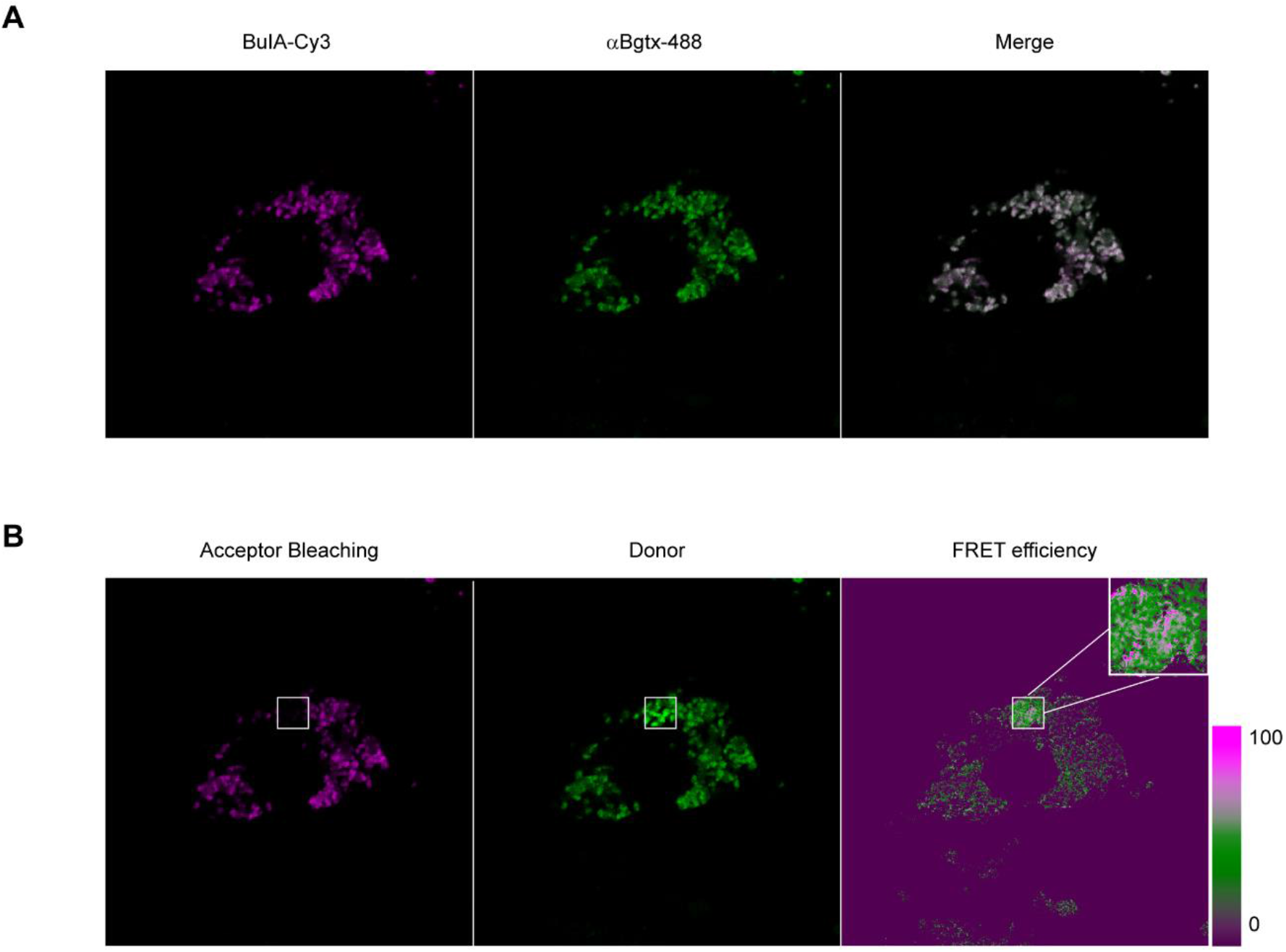
α7 and α3β4 receptors interact physically with each other. A) Original fluorescent images of human chromaffin cells treated with the β4* and the α7 receptor markers BuIA and α-BgTx, respectively. B) FRET assay, where a region of interest (ROI) of the acceptor is bleached and measured again in the donor channel. The efficiency of FRET is represented in the third panel, with the ROI enlarged.

In addition, the expression of both receptor subtypes was analyzed under various conditions: after the potentiation protocol (7 successive pulses of ACh), and after different treatments were applied to the cells prior to the potentiation protocol. These treatments included the application of three pulses of choline to desensitize the α7 receptor, the α7 receptor selective blocker ArIB, the α3β4 receptor selective blocker TxID, or ArIB and TxID simultaneously. Original fluorescence images are shown in Fig. 5A and the summary of the integrated density is plotted in Fig. 5B and C, respectively. These experiments revealed that successive ACh pulses increase α7 and α3β4 receptors expression by 1.6-fold (n=31, *p<0.05) and 1.8-fold (n=35, **p<0.01), respectively. α7, and surprisingly, also α3β4 receptor expression increases were abolished when α7 receptors were desensitized (n=11, #p<0.05, with respect to ACh receptor potentiated currents), or cells were treated with the α7 receptor antagonist ArIB (n=18, ###p<0.001 and ##p<0.01, respectively). This reflects that α7 receptors must be in an active, non-desensitized state to contribute to their own increased expression and also to the expression of the α3β4 receptor subtype. α7 receptors increased expression was also surprisingly prevented by the treatment of the cells with the α3β4 receptor blocker TxID (n=18, ###p<0.001). The increase expression of α3β4 receptors was also abolished by the treatment with TxID (n=18, ##p<0.01). When cells were treated with both toxins simultaneously, α7 and α3β4 receptors expression were not increased (n=18, ####p<0.0001 and ###p<0.001, respectively). They were even decreased with respect to control conditions in the case of α7 receptors (n=18, *p<0.05, respectively). This might reflect that part of the α7 receptors were internalized due to α7 and α3β4 receptors block.

**Figure 5.**
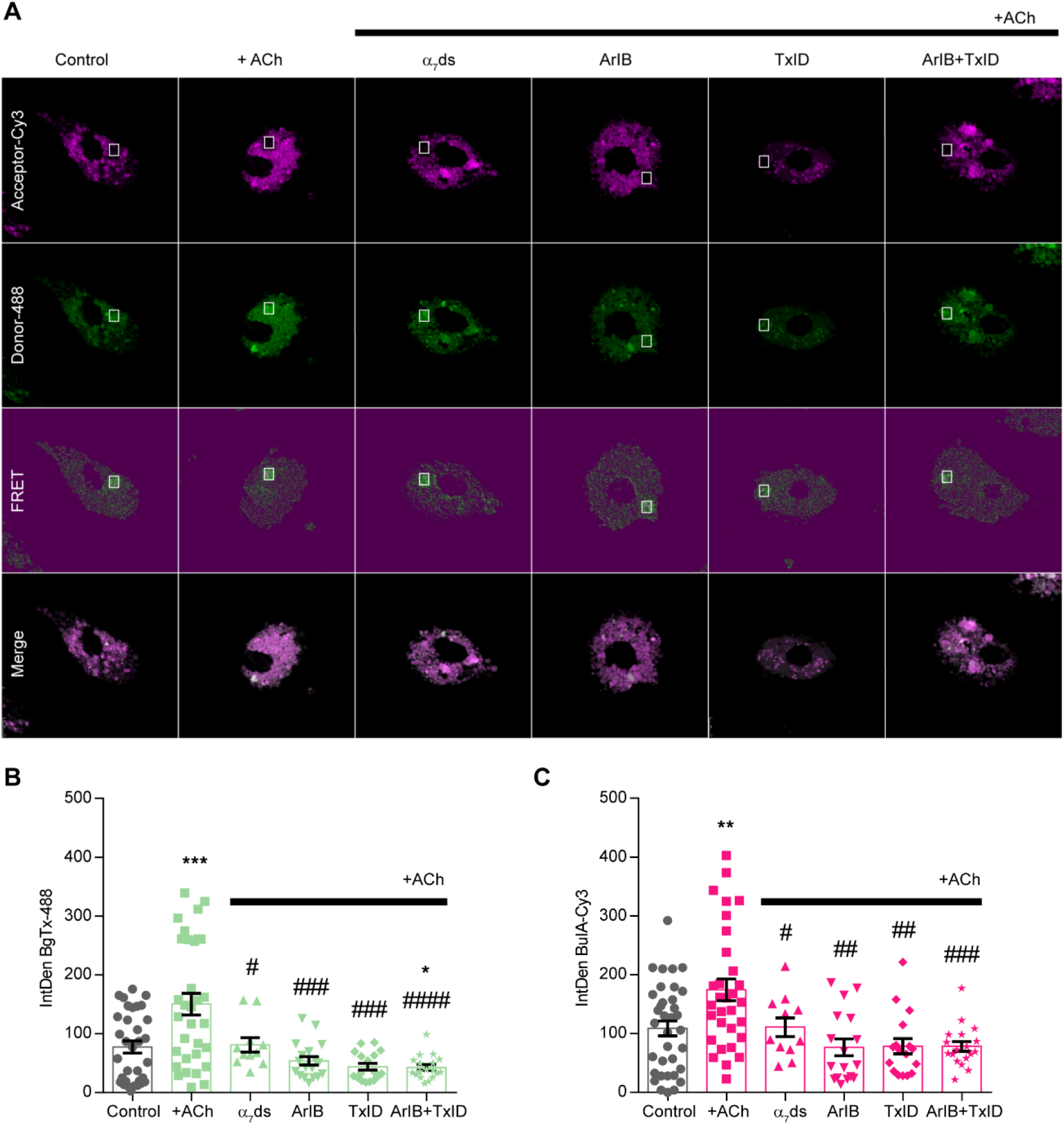
α7 and α3β4 receptors required to be in an activated state. A) Representative fluorescent images obtained under different conditions: Control; after the potentiation protocol (+ACh); after desensitization of the α7 receptor by applying 3 choline (3 mM) pulses every 90 s previously to the application of the potentiation protocol (α_7_ds); after perfusion of 100 nM ArIB (ArIB), 300 nM TxID (TxID), or ArIB and TxID 5 min before and during the application of the potentiation protocol. Cells were treated with BuIA-Cy3 (acceptor) and BgTx-488 (donor) as explained in Methods and Fig 4. B and C) Plotted data of the integrated density of BgTx-488 or BuIA-Cy3, respectively (n=18-35). Unpaired t-test. * Differences with respect to Control. # Differences with respect to +ACh.

The interaction between α7 and α3β4 was also analyzed under these conditions, giving as a result in all cases the same FRET efficiency (22.5-27.2%, n=5). Accordingly, the Correlation index values obtained for the colocalization between α7 and α3β4 receptors varied between 0.76 and 0.85 under the different conditions (n=11-35) (Table 1).

**Table 1.**
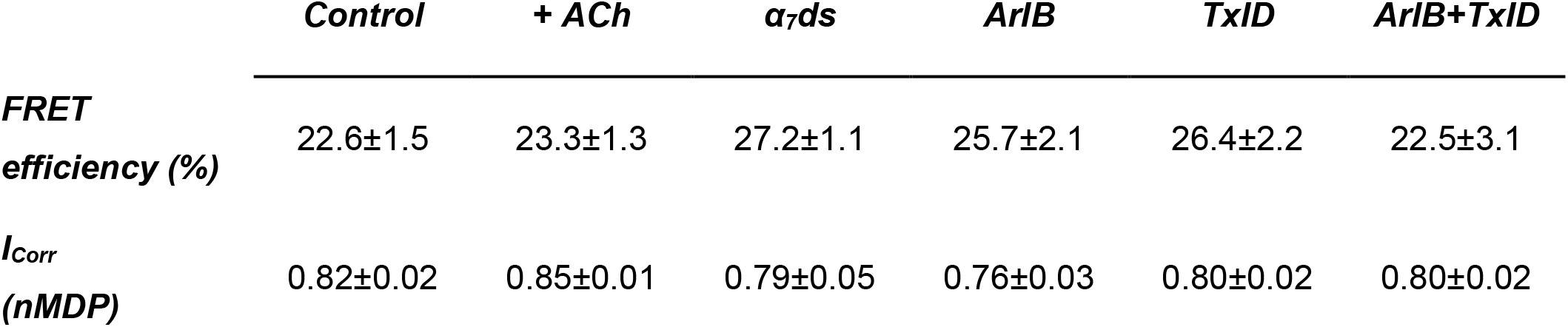
Interaction between α7 and α3β4 receptor subtypes. FRET efficacies and Correlation index (I_corr_) values obtained under different conditions of Figure 5: Control, after the potentiation protocol was applied (+ACh), or three pulses of choline every 90 s (α7ds), ArIB, TxID or ArIB plus TxID (perfused 5 min before ACh and then during the whole experiment) were applied before the potentiation protocol.

### The presence of a linked α7-α3β4 receptor complex is further supported by the toxin block of ACh and choline-evoked currents and exocytosis

The effect of the toxins ArIB and TxID, selective blockers of α7 and α3β4 receptor subtypes, respectively, might help to further understand the linkage of the nicotinic receptors in these cells. Representative recordings of the effect of ArIB and TxID on the ACh and choline-elicited currents are shown in Fig. 6A and C, for ACh and choline, respectively. The perfusion of ArIB on the ACh and choline currents diminished those currents, which partially recovered after wash-out. Then, TxID was perfused, and currents were blocked again. Dot plots of the effect of both toxins on the ACh and choline-elicited currents are shown in Fig. 6B and D, respectively. ArIB and TxID blocked the ACh-elicited currents by 52.7±9.1% (**p<0.01, n=15) and 87.6±3.1% (**p<0.01, n=8), respectively. In the case of the currents evoked by choline, ArIB and TxID block was 85±4.8% (***p<0.001, n=15) and 79.4±16.5% (*p<0.05, n=8), respectively. This means that around 90% of the nicotinic current is mediated by the α3β4 subtype (given by the TxID block of ACh currents), and 50% of those receptors are interacting with the α7 receptor subtype (given the ArIB block of ACh currents). When TxID was applied directly on the ACh or choline-elicited currents, the block achieved was 90% and 98%, respectively (n=3) (Fig. 6E and F). This means that 90% of the receptors stimulated by choline contains the α7 subtype (given by the ArIB block of choline currents), interacting all of them with the α3β4 subtype (given by the TxID block of choline currents). Thus, most of the current is carried by the α3β4 subtype, and about half of the receptors interact with the α7 subtype. Instead, all α7 receptors interact with the α3β4 subtype.

**Figure 6.**
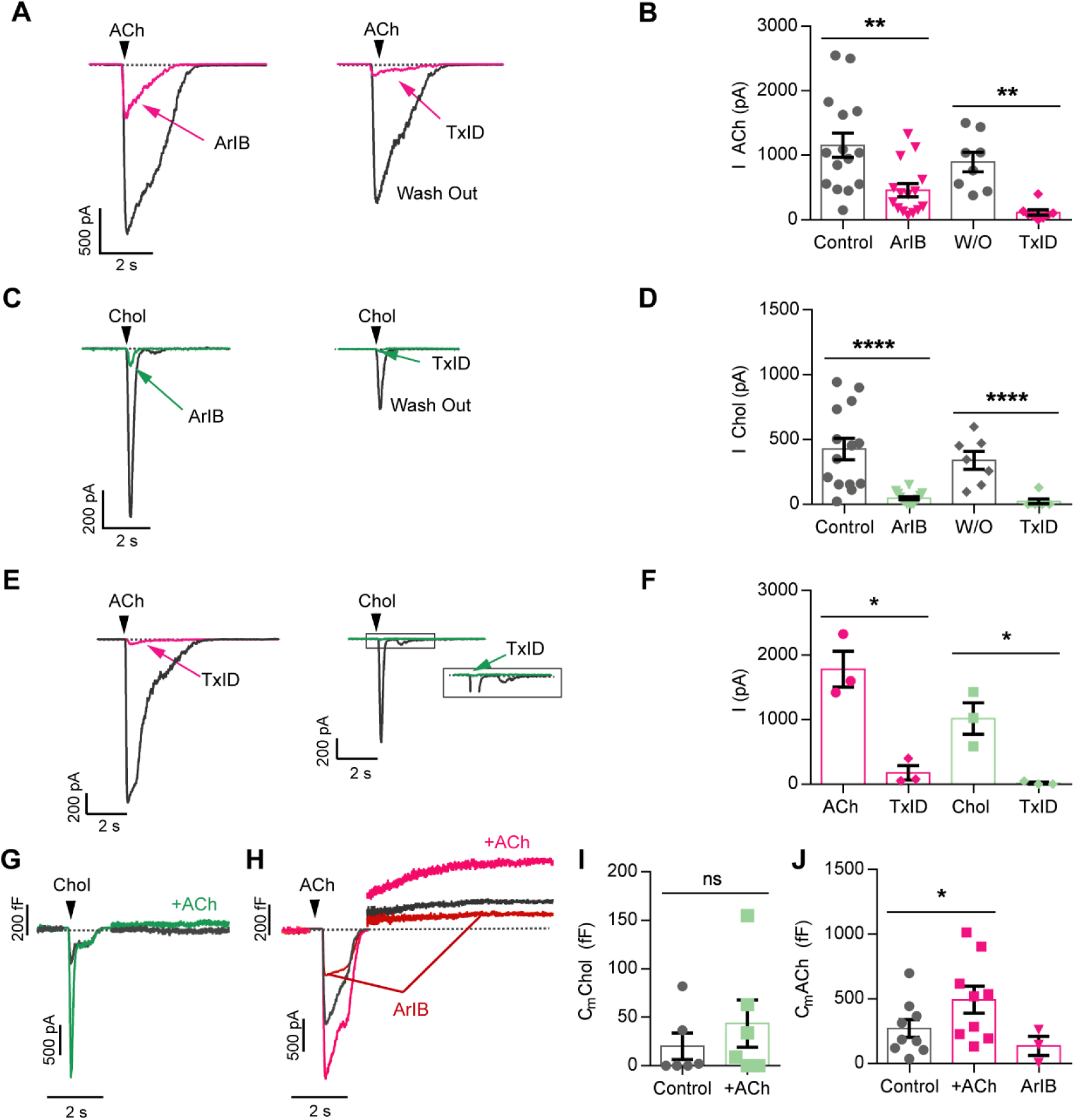
The presence of a linked α7-α3β4 receptor complex is further supported by the toxin block of ACh and choline-evoked currents and exocytosis. Representative recordings of the nicotinic currents elicited by ACh (A) or choline (C) obtained after perfusion with ArIB or TxID, and their corresponding wash-out. The peak values of the currents elicited by ACh or Choline are summarized in panels B (n=8-15) and D (n=8-15). E and F) Block of currents evoked by ACh or choline by TxID, and the corresponding plotted data showing the summary of the peak current values obtained with both stimuli. G) Nicotinic currents and plasma membrane capacitance traces elicited by choline before and after application of 7 pulses of ACh to obtain the α7 receptor potentiated currents (+ACh). H) Nicotinic currents and C_m_ traces elicited by ACh before and after application of the potentiation protocol to obtain the receptor potentiated currents (+ACh). Then, ArIB at 100 nM concentration was perfused for 5 min before the pulse, and currents and C_m_ were inhibited by this toxin (n=3-9). Paired t-test. I and J) Average C_m_ values corresponding to the experiments of panel G and H are drawn in I (n=6), and J (n=2-9), respectively. Paired t-test.

In a different set of experiments, toxins were tested on exocytosis, determined by C_m_ measurements. A single pulse of 500 ms of choline was applied every 3 min. Choline did not evoke exocytosis due to calcium entry through the nicotinic receptor (20.1±13.7 fF) (n=6). After the potentiation protocol was applied, the increased choline currents elicited 43.5±24.4 fF (n=6), which did not exhibit significant statistical differences with respect to control exocytosis (Fig.6G). Thus, the nicotinic current that flows through the α7 receptor subtype did not evoke exocytosis. However, in different cells, a single pulse of 500 ms ACh was applied, eliciting a large exocytosis that amounted to 271.3±69.3 fF (n=9). Then the potentiation protocol was applied to obtain the receptor potentiated currents. The exocytosis triggered amounted to 492.8±104.9 fF (n=9, *p<0.05). Afterwards, ArIB was perfused and the exocytosis decreased (137.2±74.2 fF) (n=3) (Fig. 6H). Data are summarized in the dot plots of Fig. 6I and J. These experiments show that the α7 receptor does contribute to exocytosis only when the α3β4 receptor is fully activated by ACh, and not when it is partially activated by choline. The α3β4 receptor instead is able to evoke exocytosis in the absence of α7 receptor activation, probably due to the receptor pool which does not interact with the α7 receptor subtype.

### Age dependence of the interaction and sex dependence of the expression of α7 and α3β4 receptor subtypes

The degree of physical interaction between α7 and α3β4 receptor subtypes was investigated by comparing FRET efficiency in donors sorted in two groups of age, below and above 50 years old. BuIA-Cy3 and BgTx-488 were used as acceptor and donor, respectively, in the FRET experiments. Original images are shown in Fig. 7A. A dot plot of the average FRET efficiency obtained under both conditions are shown in Fig. 7B. FRET efficiency values were 30.6±2.7 (n=8 donors) for the group of less than 50 years old and 21.6±1.5 (n=27 donors) for the group of more than 50 years old. Significant statistical differences were obtained between both groups (**p<0.01). This means that the physical interaction between α7 and α3β4 receptor subtypes decreased above 50 years old. However, expression of both receptor subtypes did not vary with age (Fig. 7C). A decrease in the efficiency of the interaction is expected to produce an increase in the degree of the α7 receptor desensitization by choline (protocol of Fig. 2B). And this was the case. The percentage of desensitization of ACh was lower in the group of age below 50 years old (10.7±8.4%, n=8 donors), with respect to the group of more than 50 years old (45.3±5.8%, n=7 donors) (**p<0.01). Representative ACh-elicited currents and plotted data are shown in Fig. 7D and E, respectively. Thus, the physical interaction between α7 and α3β4 receptor subtypes decreases with age, increasing the desensitization rate.

**Figure 7.**
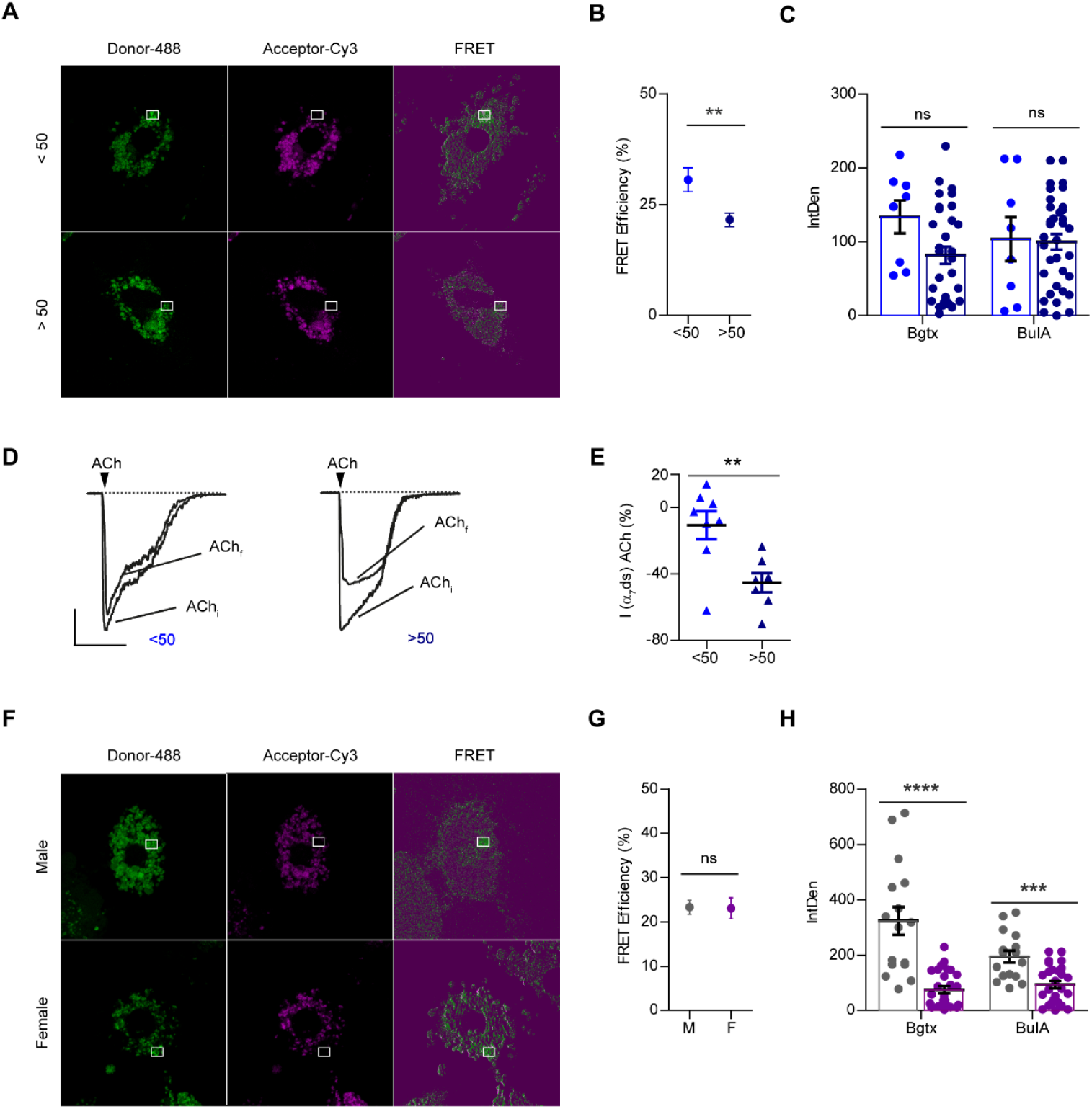
Age and sex dependence of the interaction and expression of α7 and α3β4 receptor subtypes. A) Representative FRET images using BuIA-Cy3 as acceptor and BgTx-488 as donor. Donors were sorted in two groups of age, below and above 50 years old. B) The percentage of FRET efficiency is averaged in both groups of age, below (light blue) and above (dark blue) 50 years old (n donors <50 years old=8, n donors >50 years old=27). Unpaired t-test. C) Dot plot that summarizes the expression data of both groups of age, below (light blue) and above (dark blue) 50 years old. D) Representative recordings of the first (ACh_i_) and the last (ACh_f_) nicotinic currents elicited by ACh of a set of 7 pulses obtained in cells from donors of age below and above 50 years old. Three pulses of choline were applied at the beginning of the experiment, before the ACh pulses were applied (protocol of Fig. 2B). E) The percentage of desensitization of the currents calculated as the percentage of decay of the last ACh-nicotinic peak current with respect to the first one is plotted in both groups of age, below (light blue) and above (dark blue) 50 years old (n donors <50 years old=8, n donors >50 years old=7). Unpaired t-test. F) FRET images obtained using BuIA-Cy3 as acceptor and BgTx-488 as donor in male and female adrenal gland donors. G) The percentage of FRET efficiency is averaged in male (M) and females (F). H) Dot plot that summarizes the expression data of α7 and α3β4 receptors in panel F calculated as integrated density of BgTx-488 or BuIA-Cy3, respectively, in males (grey) and females (violet) (n=16 cells from men, 27 cells from women). BgTx: U Mann Whitney; BuIA: Unpaired t-test.

With respect to sex, FRET efficiency did not vary between males (23.3±1.6, n=4 donors) and females (23.1±2.4, n=6 donors). Representative images and data graph are shown in Fig. 7F and G, respectively. However, expression of α7 and α3β4 receptor subtypes was larger in men with respect to women (Fig 7H). Total BgTx fluorescence amounted to 323.5±50.5 (n=16) in males, and to 75±12.6 (n=27) in females (****p<0.0001). Also, total BuIA fluorescence amounted to 194.7±21.5 (n=16) in males, and to 93.3±13 (n=27) in females (***p<0.001). Therefore, males possess a significant larger number of α7 (4-fold) and α3β4 (2-fold) nicotinic receptors than females.

## Discussion

Much of our current understanding regarding the functionality of the α7 receptor derives from studies performed in recombinant receptors reconstituted in heterologous systems or non-human native cells. However, the α7 receptor subtype is always expressed together other non-α7 receptor subtypes in the different tissues. The goal of the present study was to investigate physical and/or functional interactions between native human α7 and non-α7 nicotinic receptor subtypes, their calcium regulation and their age and sex dependence. In this study we have used a native human postganglionic sympathetic neuron, the chromaffin cell of the adrenal gland. It is the major adrenaline-secreting cell in our organism. This cell mainly expresses α7 and α3β4 receptor subtypes (Hone *et al.*, 2015; Hone *et al.*, 2017; Perez-Alvarez *et al.*, 2012a; Perez-Alvarez *et al.*, 2012b). Here we show that these receptor subtypes physically interact with each other in human chromaffin cells. Interaction between native nicotinic receptor subtypes has not previously been reported. However, in *Xenopus oocytes*, Criado and colleagues (Criado *et al.*, 2012) showed that stable and unstable interactions between α7 with β4 or α3 subunits, respectively, were established among human and bovine receptor subunits. Our data indicate that α7 and α3β4 receptors exist as separate entities and interact with each other forming a receptor complex. The following evidences support this conclusion: 1) Interaction between subunits determined by FRET (Fig. 4B); 2) desensitization of the α7 subtype independently of the α3β4 subtype receptor (Fig. 2B); 3) block of currents and exocytosis elicited by ACh or choline by selective α7 and α3β4 receptor blockers (Fig. 5). Furthermore, the interaction between α7 and α3β4 receptor subtypes makes possible that each receptor subtype controls the functionality and expression of the other subtype. This interaction takes place even if the receptors are blocked or desensitized.

The desensitization of the α7 receptor was prevented by full α3β4 receptor activation with ACh, but not by partial activation with choline. This is an extremely relevant finding of the present study which shows that in vivo, after ACh stimulation, α7 receptors do not desensitize due to the interaction with fully activated α3β4 receptors. This allows the cell to largely increase nicotinic currents and exocytosis, and therefore, catecholamine release. Our data put the focus on α3β4 receptors as pharmacological targets to improve α7 receptors activity.

On the other hand, choline, which is obtained after degradation of ACh by the acetylcholinesterase, is able to desensitize the α7 receptor due to its high affinity for this subtype and its low affinity for the α3β4 receptor subtype. As α3β4 receptors are not fully activated, α7 subtypes can be desensitized after repetitive choline pulses. In this way choline acts as a brake to avoid the uncontrolled potentiation of nicotinic currents, exhibiting a prominent role in the regulation of the activity of nicotinic receptors.

It is relevant to note that the nicotinic currents triggered by choline, both control or “potentiated”, were not able to evoke exocytosis. This was due to a partial but not full activation of the α3β4 receptors by choline. In the case of ACh, full agonist of both receptor subtypes, exocytosis was triggered. In addition, the selective α7 receptor blocker ArIB was able to diminish the exocytosis elicited by ACh, indicating that α7 receptors, in spite of not triggering exocytosis, cooperate with the normal functioning of α3β4 receptors. This reflects once more that both receptors work together, and if one of them is affected, the activity of the other one is also modified.

In relation with the subunit receptor composition, the use of ArIB and TxID might help to clarify it. ArIB blocked half of the ACh-triggered current, and TxID around 90% of the total current, which means that only a sub-population of the α3β4 receptors is bound to α7 receptors. However, ArIB and TxID block of choline-elicited currents amounted to around 90%, which reflects that most α7 receptors are bound to α3β4 receptors.

We have also found that the physical interaction between α7 and α3β4 receptors decreased in donors with ages above 50 years old. The decrease of the efficiency of the interaction between nicotinic receptors provoked an increased α7 receptor desensitization of both receptor subtypes and a decrease of the nicotinic currents, impairing the cholinergic function with age. This is a very important finding that may help to explain the functional decline of the cholinergic activity in normal aging, in addition to other mechanisms (Schliebs and Arendt, 2011). The fact that the present study is performed in human cells gives even more relevance to that finding. In the literature it has been also reported that in humans, there is an age-associated diminished expression of high-affinity [3H]nicotine ligand binding sites (Giacobini, 1992; Kellar *et al.*, 1987; Schröder *et al.*, 1991; Whitehouse *et al.*, 1988; Whitehouse *et al.*, 1986). This also occurs in the rodent brain (Araujo *et al.*, 1990; Schulz *et al.*, 1993; Zhang *et al.*, 1990).

In addition, our data shows that expression, but not interaction, of α7 and α3β4 nicotinic receptors depends on sex. Men exhibited 2-3-fold increase of BgTx and BuIA receptors labeling with respect to women. It might be that the direct influence of adrenal gland steroids regulates expression of nicotinic receptors in the chromaffin cell. Indeed, estradiol increases α7 receptor subunit expression in the dorsal raphe and locus coeruleus of macaques (Centeno *et al.*, 2006) and BgTx binding in the suprachiasmatic nucleus (Miller *et al.*, 1984), and also potentiates human α4β2 receptors (Curtis *et al.*, 2002; Jin & Steinbach, 2015; Paradiso *et al.*, 2000; Paradiso *et al.*, 2001). Recombinant human α3β4 receptors are noncompetitively inhibited by estrogens (Ke & Lukas, 1996; Nakazawa & Ohno, 2001), but without significant effects on the number of receptor radioligand-binding sites (Ke & Lukas, 1996). Our data constitute the first study in which human native α7 and α3β4 nicotinic receptors have been determined. If this ratio in the expression of receptors is maintained in the brain, it might help to explain sex-dependent incidence differences of diseases such as Alzheimer’s disease. However, our present knowledge in rodent brain indicates that male exhibits lower amount (Chiari *et al.*, 1999; Koylu *et al.*, 1997) or no differences (Cosgrove *et al.*, 2012) of β2-containing nicotinic receptors with respect to female rats. It could be that the variations between the sexes in the human brain are very different from what occurs in non-human species.

## Materials and Methods

### Reagents

Acetylcholine chloride, choline chloride, amphotericin B, penicillin/streptomycin, protease type XIV, collagenase type I, poly-D-lysine hydrobromide, red blood cell lysis solution, dimethylsulfoxide, bovine serum albumin (BSA), aprotinin, leupeptin trifluoroacetate, pepstatin A, 1,1-Dimethyl-4-phenylpiperazinium iodide (DMPP), PNU-282987 hydrate and all salts were purchased from Merk Sigma-Aldrich (St. louis, MO, USA). Dulbecco’s Modified Eagle’s Medium (DMEM), Glutamax, α-Bungarotoxin (BgTx) Alexa Fluor 488 conjugate, ProLong Gold Antifade Mountant were purchased from ThermoFisher Scientific (Massachusetts, USA). Fetal bovine serum was obtained from LabClinics (Barcelona, Spain).

### Peptide synthesis

The α-conotoxin (α-Ctx) ArIB (V11,V16D) (ArIB), was synthesized as previously described (Whiteaker *et al.*, 2007). The α-Ctx TxID (TxID) (Luo *et al.*, 2013) was synthesized using an Apex 396 automated peptide synthesizer (AAPPTec, Louisville, KY) according to previously described methods (Hone *et al.*, 2013).

### Cell culture of human chromaffin cells

Human adrenal chromaffin cells were isolated and cultured as previously reported (Hone *et al.*, 2015). Briefly, glands were perfused through the vein with protease type XIV and after dissecting the gland, the medullary tissue was extracted and digested by incubation with collagenase type I. Cells were isolated after several filtrations and plated on poli-L-lysine (0.1 mg/ml) coated coverslips. Cells were maintained in DMEM medium supplemented with 1% Glutamax, 10% fetal bovine serum (FBS) and penicillin/streptomycin, at 37° C in an incubator under an atmosphere of 95% air and 5% CO2 for up to 7 days. The culture medium was changed daily by exchanging approximately 70% of the solution with fresh medium.

### Electrophysiological recordings of “patch-clamp”

To perform the recordings of nicotinic currents in the perforated-patch configuration, the extracellular solution was (in mM): 2 CaCl_2_, 145 NaCl, 5 KCl, 1 MgCl_2_, 10 HEPES and 10 glucose (pH 7.4; 315 mOsM). When extracellular free-calcium solutions were used, 5 mM EGTA was added to this solution in the absence of calcium. The composition of the intracellular solution was (in mM): 145 Kglutamate, 8 NaCl, 1 MgCl2, 10 HEPES and 0.5 Amphotericin B (pH 7.2; 322 mOsM). ACh was applied at 300 μM. An amphotericin B stock solution was prepared every day at a concentration of 50 mg/mL in dimethyl sulphoxide (DMSO) and kept protected from light. The final concentration of amphotericin B was prepared by ultrasonicating 10 μL of stock amphotericin B in 1 mL of internal solution in the darkness. Pipettes were tip-dipped in amphotericin-free solution for several seconds and back-filled with freshly mixed intracellular amphotericin solution.

The electrophysiological measurements were carried out using an EPC-10 USB amplifier and the PatchMaster program (HEKA Elektronik, Germany). 2-3 MΩ borosilicate glass polished pipettes were used. After the formation of the seal and the perforation, only those recordings with a leakage current below 20 pA were accepted. The series resistant was 17.7±3 MΩ and 16.3±2 MΩ at the beginning and at the end of the experiment, respectively (n=28). This resistance was compensated electronically up to 70% in the voltage-clamp mode. Cell membrane capacitance (C_m_) changes as an index of exocytosis were estimated by the Lindau-Neher technique implemented in the “Sine+DC” feature of the PatchMaster lock-in software. A 1 kHz, 70 mV peak-to-peak amplitude sinewave was applied at a holding potential (V_h_) of −80 mV.

To conduct the experiments, cells were gravity perfused with the extracellular solution delivered through a polyethylene tube placed close to the cell of interest. To ensure rapid solution exchange, a multi barrel pipette was used for agonist delivery. Agonist pulses (500 ms pulses) were applied by means of a valve controller triggered by the amplifier. The extracellular solution without the agonist was immediately perfused once the agonist application ended. The fluid level was continuously controlled by a home-made fibre optics system coupled to a pump that removed excess fluid.

A “potentiation protocol” was performed in the electrophysiological experiments consisting of the application of successive ACh pulses every 90 s until the current reached the maximal stable amplitude.

The analysis of data was done using IGOR Pro software (WaveMetrics, OR).

### Cell treatments

In the electrophysiological recordings and image experiments, ArIB and TxID were perfused at 100 nM and 300 nM, respectively, during 5 minutes before the start of the protocol and throughout the remainder of the experiment.

### Fluorescence

Cells plated on 12 mm glass coverslips were quick rinsed with PBS and then fixed with 2% paraformaldehyde (PFA) for 20 min. After that, three washing steps of 5 min with PBS were performed. Incubation of fluorescent labeled toxins was performed for 30-40 min on ice in a buffer containing 0.1 mg/ml bovine serum albumin (BSA) to block non-specific binding and 10 μg/ml each of aprotinin, leupeptin trifluoroacetate, and pepstatin A to prevent peptide degradation by proteases. This was followed by extensive wash-out with PBS. Finally, coverslips were mounted in ProLong.

To study the interaction and expression of α3β4 and α7 receptor subtypes in human cells, α-Ctx BuIA labeled with Cy3 (BuIA-Cy3) (300 nM) and α-Bungarotoxin (BgTx) labeled with Alexa 488 (BgTx-488) (1.2 μM) were used, respectively. To define non-specific binding, control coverslips were treated with 2 nM nicotine 15 min before and during the fluorescent toxins incubation. As a negative control, some coverslips were labeled with only one fluorescent toxin. As an additional negative control, FRET was also performed in non-chromaffin cells of the same coverslip.

A “potentiation protocol” was performed in the expression experiments consisting of the application of 7 successive ACh pulses every 90 s.

### Microscope image acquisition and analysis

Microscopy images were collected using a spectral confocal microscope (Leica TCS SP5) equipped with argon laser (wavelength 488 nm), DPSS laser (wavelength 561 nm) and helium-neon laser (wavelengths 594 nm and 633 nm) for the excitation of fluorescent probes and a 63X/ 1.4-0.6 oil immersion objective. Images were acquired in a medial plane in each cell under identical conditions to facilitate statistical analysis across conditions.

Expression of the different nicotinic receptor subtypes was determined by measuring the integrated density (IntDen), which is the product of the area and the mean gray value, by using Fiji-ImageJ software. Also, the Fiji-ImageJ colocalization Colormap plugin was used to calculate the colocalization Correlation index values (I_Corr_). To perform and analyze FRET experiments, wizard Acceptor Photobleaching (AB) (LAS AF, Leica) was used. Only those measurements where the acceptor was bleached 80% or more were accepted. Red images were converted to magenta, to ensure legibility for color-blind people.

### Statistics

Data are given as mean±SEM. A Kolmogorov-Smirnov Normality test was first performed. Then, t-Student, U Mann Whitney o Wilcoxon signed rank tests were used. Statistical analysis was performed using IBM SPSS Statistics Version 24.0. (Armonk, NY). The number of data n refers to number of cells unless otherwise stated. Differences were accepted as significantly different when the p-value was under 0.05 (plot with an asterisk *), 0.01 (**), 0.001 (***) and 0.0001 (****).

### Study approval

The human adrenal glands used in this study were collected from 30 organ donors in three hospitals in Madrid (12 de Octubre, Fundación Jiménez Díaz and Clínico San Carlos). Written consents were signed by donor’s relatives. The protocol was approved by the Ethics Committee of Autonomous University of Madrid and by review boards of each hospital.

## Acknowledgments

This work was supported by grants from the Spanish Ministerio de Ciencia y Tecnología (BFU2012-30997 and BFU2015-69092 to A.A.) and the U.S. National Institutes of Health (GM136430 and GM103801 to J.M.M).

## Author Contributions

Participated in research design: A.A., A.J.P. Conducted experiments: A.J.P., S.S.L. Performed data analysis: A.J.P. Contributed human adrenal glands: J.M.P, C.G.E., J.B. Contributed fluorescent toxins: J.M.M. Wrote the manuscript: A.A., A.J.P.

## References

Albillos, A., Dernick, G., Horstmann, H., Almers, W., De Toledo, G. A., & Lindau, M. (1997). The exocytotic event in chromaffin cells revealed by patch amperometry. Nature, 389(6650), 509–512.

Araujo, D. M., Lapchak, P. A., Meaney, M. J., Collier, B., & Quirion, R. (1990). Effects of aging on nicotinic and muscarinic autoreceptor function in the rat brain: Relationship to presynaptic cholinergic markers and binding sites. The Journal of Neuroscience: The Official Journal of the Society for Neuroscience, 10(9), 3069–3078.

Centeno, M. L., Henderson, J. A., Pau, K. F., & Bethea, C. L. (2006). Estradiol increases α7 nicotinic receptor in serotonergic dorsal raphe and noradrenergic locus coeruleus neurons of macaques. Journal of Comparative Neurology, 497(3), 489–501.

Chiari, A., Tobin, J. R., Pan, H., Hood, D. D., & Eisenach, J. C. (1999). Sex differences in cholinergic analgesia I a supplemental nicotinic mechanism in normal females. Anesthesiology: The Journal of the American Society of Anesthesiologists, 91(5), 1447.

Cosgrove, K. P., Esterlis, I., McKee, S. A., Bois, F., Seibyl, J. P., Mazure, C. M.,… O’Malley, S. S. (2012). Sex differences in availability of β2*-nicotinic acetylcholine receptors in recently abstinent tobacco smokers American Medical Association.

Criado, M., Valor, L. M., Mulet, J., Gerber, S., Sala, S., & Sala, F. (2012). Expression and functional properties of alpha7 acetylcholine nicotinic receptors are modified in the presence of other receptor subunits. Journal of Neurochemistry, 123(4), 504–514. doi:10.1111/j.1471-4159.2012.07931.x [doi]

Curtis, L., Buisson, B., Bertrand, S., & Bertrand, D. (2002). Potentiation of human α4β2 neuronal nicotinic acetylcholine receptor by estradiol. Molecular Pharmacology, 61(1), 127–135.

Giacobini, E. (1992). Nicotine acetylcholine receptors in human cortex: Aging and Alzheimer’s disease. Biology of Nicotine, 183–215.

Hone, A. J., McIntosh, J. M., Azam, L., Lindstrom, J., Lucero, L., Whiteaker, P.,… Albillos, A. (2015). Alpha-conotoxins identify the alpha3beta4* subtype as the predominant nicotinic acetylcholine receptor expressed in human adrenal chromaffin cells. Molecular Pharmacology, 88(5), 881–893. doi:10.1124/mol.115.100982 [doi]

Hone, A. J., Michael McIntosh, J., Rueda-Ruzafa, L., Passas, J., de Castro-Guerín, C., Blázquez, J.,… Albillos, A. (2017). Therapeutic concentrations of varenicline in the presence of nicotine increase action potential firing in human adrenal chromaffin cells. Journal of Neurochemistry, 140(1), 37–52.

Hone, A. J., Ruiz, M., Christensen, S., Gajewiak, J., Azam, L., & McIntosh, J. M. (2013). Positional scanning mutagenesis of α-conotoxin PeIA identifies critical residues that confer potency and selectivity for α6/α3β2β3 and α3β2 nicotinic acetylcholine receptors. Journal of Biological Chemistry, 288(35), 25428–25439.

Jin, X., & Steinbach, J. H. (2015). Potentiation of neuronal nicotinic receptors by 17β-estradiol: Roles of the carboxy-terminal and the amino-terminal extracellular domains. PLoS One, 10(12), e0144631.

Ke, L., & Lukas, R. J. (1996). Effects of steroid exposure on ligand binding and functional activities of diverse nicotinic acetylcholine receptor subtypes. Journal of Neurochemistry, 67(3), 1100–1112.

Kellar, K. J., Whitehouse, P. J., Martino-Barrows, A. M., Marcus, K., & Price, D. L. (1987). Muscarinic and nicotinic cholinergic binding sites in Alzheimer’s disease cerebral cortex. Brain Research, 436(1), 62–68.

Koylu, E., Demirgören, S., London, E. D., & Pöğün, S. (1997). Sex difference in up-regulation of nicotinic acetylcholine receptors in rat brain. Life Sciences, 61(12), PL185–PL190.

Luo, S., Zhangsun, D., Zhu, X., Wu, Y., Hu, Y., Christensen, S.,… McIntosh, J. M. (2013). Characterization of a novel alpha-conotoxin TxID from conus textile that potently blocks rat alpha3beta4 nicotinic acetylcholine receptors. Journal of Medicinal Chemistry, 56(23), 9655–9663. doi:10.1021/jm401254c [doi]

Miller, M. M., Silver, J., & Billiar, R. B. (1984). Effects of gonadal steroids on the in vivo binding of [125I] α-bungarotoxin to the suprachiasmatic nucleus. Brain Research, 290(1), 67–75.

Nakazawa, K., & Ohno, Y. (2001). Modulation by estrogens and xenoestrogens of recombinant human neuronal nicotinic receptors. European Journal of Pharmacology, 430(2-3), 175–183.

Paradiso, K., Sabey, K., Evers, A. S., Zorumski, C. F., Covey, D. F., & Steinbach, J. H. (2000). Steroid inhibition of rat neuronal nicotinic α4β2 receptors expressed in HEK 293 cells. Molecular Pharmacology, 58(2), 341–351.

Paradiso, K., Zhang, J., & Steinbach, J. H. (2001). The C terminus of the human nicotinic α4β2 receptor forms a binding site required for potentiation by an estrogenic steroid. Journal of Neuroscience, 21(17), 6561–6568.

Pérez-Alvarez, A., & Albillos, A. (2007). Key role of the nicotinic receptor in neurotransmitter exocytosis in human chromaffin cells. Journal of Neurochemistry, 103(6), 2281–2290. doi: 10.1111/j.1471-4159.2007.04932.x

Perez-Alvarez, A., Hernandez-Vivanco, A., Alonso Y Gregorio, S., Tabernero, A., McIntosh, J. M., & Albillos, A. (2012a). Pharmacological characterization of native alpha7 nicotinic ACh receptors and their contribution to depolarization-elicited exocytosis in human chromaffin cells. British Journal of Pharmacology, 165(4), 908–921. doi:10.1111/j.1476-5381.2011.01596.x [doi]

Perez-Alvarez, A., Hernandez-Vivanco, A., McIntosh, J. M., & Albillos, A. (2012b). Native alpha6beta4* nicotinic receptors control exocytosis in human chromaffin cells of the adrenal gland. FASEB Journal: Official Publication of the Federation of American Societies for Experimental Biology, 26(1), 346–354. doi:10.1096/fj.11-190223 [doi]

Schröder, H., Giacobini, E., Struble, R. G., Luiten, P. G., van der Zee, Eddy A, Zilles, K., & Strosberg, A. (1991). Muscarinic cholinoceptive neurons in the frontal cortex in alzheimer’s disease. Brain Research Bulletin, 27(5), 631–636.

Schulz, D. W., Kuchel, G. A., & Zigmond, R. E. (1993). Decline in response to nicotine in aged rat striatum: Correlation with a decrease in a subpopulation of nicotinic receptors. Journal of Neurochemistry, 61(6), 2225–2232.

Schliebs, R.& Arendt, T. (2011) The cholinergic system in aging and neuronal degeneration (2011) Behav Brain Research 221(2), 555–63. doi: 10.1016/j.bbr.2010.11.058

Whiteaker, P., Christensen, S., Yoshikami, D., Dowell, C., Watkins, M., Gulyas, J.,… McIntosh, J. M. (2007). Discovery, synthesis, and structure activity of a highly selective α7 nicotinic acetylcholine receptor antagonist. Biochemistry, 46(22), 6628–6638.

Whitehouse, P. J., Martino, A. M., Wagster, M. V., Price, D. L., Mayeux, R., Atack, J. R., & Kellar, K. J. (1988). Reductions in [3H] nicotinic acetylcholine binding in Alzheimer’s disease and Parkinson’s disease: An autobiographic study. Neurology, 38(5), 720.

Whitehouse, P. J., Martino, A. M., Antuono, P. G., Lowenstein, P. R., Coyle, J. T., Price, D. L., & Kellar, K. J. (1986). Nicotinic acetylcholine binding sites in Alzheimer’s disease. Brain Research, 371(1), 146–151.

Zhang, X., Wahlström, G., & Nordberg, A. (1990). Influence of development and aging on nicotinic receptor subtypes in rodent brain. International Journal of Developmental Neuroscience, 8(6), 715–721.

